# Common scab disease: structural basis of elicitor recognition in pathogenic *Streptomyces* species

**DOI:** 10.1101/2023.05.10.540135

**Authors:** Frédéric Kerff, Samuel Jourdan, Isolde M. Francis, Benoit Deflandre, Silvia Ribeiro Monteiro, Nudzejma Stulanovic, Rosemary Loria, Sébastien Rigali

**Affiliations:** InBioS – Center for Protein Engineering, Institut de Chimie B6a, University of Liège, Liège, Belgium; Department of Biology, California State University, Bakersfield, CA, USA; Department of Plant Pathology, University of Florida, Gainesville, FL, USA

**Keywords:** Host-pathogen interaction, elicitor binding, sugar transport, carbohydrate metabolism, plant pathogen, ligand-protein interaction

## Abstract

In *Streptomyces scabiei*, the main causative agent of common scab disease of root and tuber crops, the interaction between the substrate-binding protein (SBP) CebE (CebE^scab^) and cellotriose released by the plant host (*K*_D_ in the nanomolar range) is the first event for the onset of its pathogenic lifestyle. Here we report the structure of CebE^scab^ in complex with cellotriose at a 1.55 Å resolution, adopting a general fold of the B subcluster of SBPs. The interaction between CebE^scab^ and cellotriose involves multiple direct or water-mediated hydrogen bonds and hydrophobic interactions, the glucose monomer at the non-reducing end occupying the most conserved part of the substrate-binding cleft. As main interactions between the two domains of CebE involve cellotriose itself, the closed conformational state of CebE is performed via an induced-fit ligand binding mechanism where cellotriose binding triggers the domain movement. Analysis of regulon predictions revealed that the signaling pathway from the CebE-mediated cellotriose transport to the transcriptional activation of thaxtomin phytotoxin biosynthesis is conserved in *Streptomyces* spp causing common scab, except for *Streptomyces ipomoeae* that specifically colonizes sweet potatoes and responds to other and yet unknown virulence elicitors. Interestingly, strains belonging to pathogenic species *turgidiscabies* and *caniscabies* have a cellotriose-binding protein orthologous to the CebE protein of the saprophytic species *Streptomyces reticuli* with lower affinity for its substrate (*K*_D_ in the micromolar range), suggesting higher cellotriose concentrations for perception of their host. Our work also provides the structural basis for the uptake of cellobiose and cellotriose by non-pathogenic cellulose-decomposing *Streptomyces* species.

**Importance:** Common scab is a disease caused by few *Streptomyces* species that affects important root and tuber crops including potato, beet, radish, and parsnip, resulting in major economic losses worldwide. In this work we unveiled the molecular basis of host recognition by these pathogens by solving the structure of the sugar-binding protein CebE of *S*. *scabiei* in complex with cellotriose, the main elicitor of the pathogenic lifestyle of these bacteria. We further revealed that the signaling pathway from CebE-mediated transport of cellotriose is conserved in all pathogenic species except *S*. *ipomoeae* that causes soft rot disease on sweet potatoes. Our work also provides the structural basis of the uptake of cellobiose and cellotriose in saprophytic *Streptomyces* species, the first step activating the expression of the enzymatic system degrading the most abundant polysaccharide on earth, cellulose.

**Graphical abstract:** 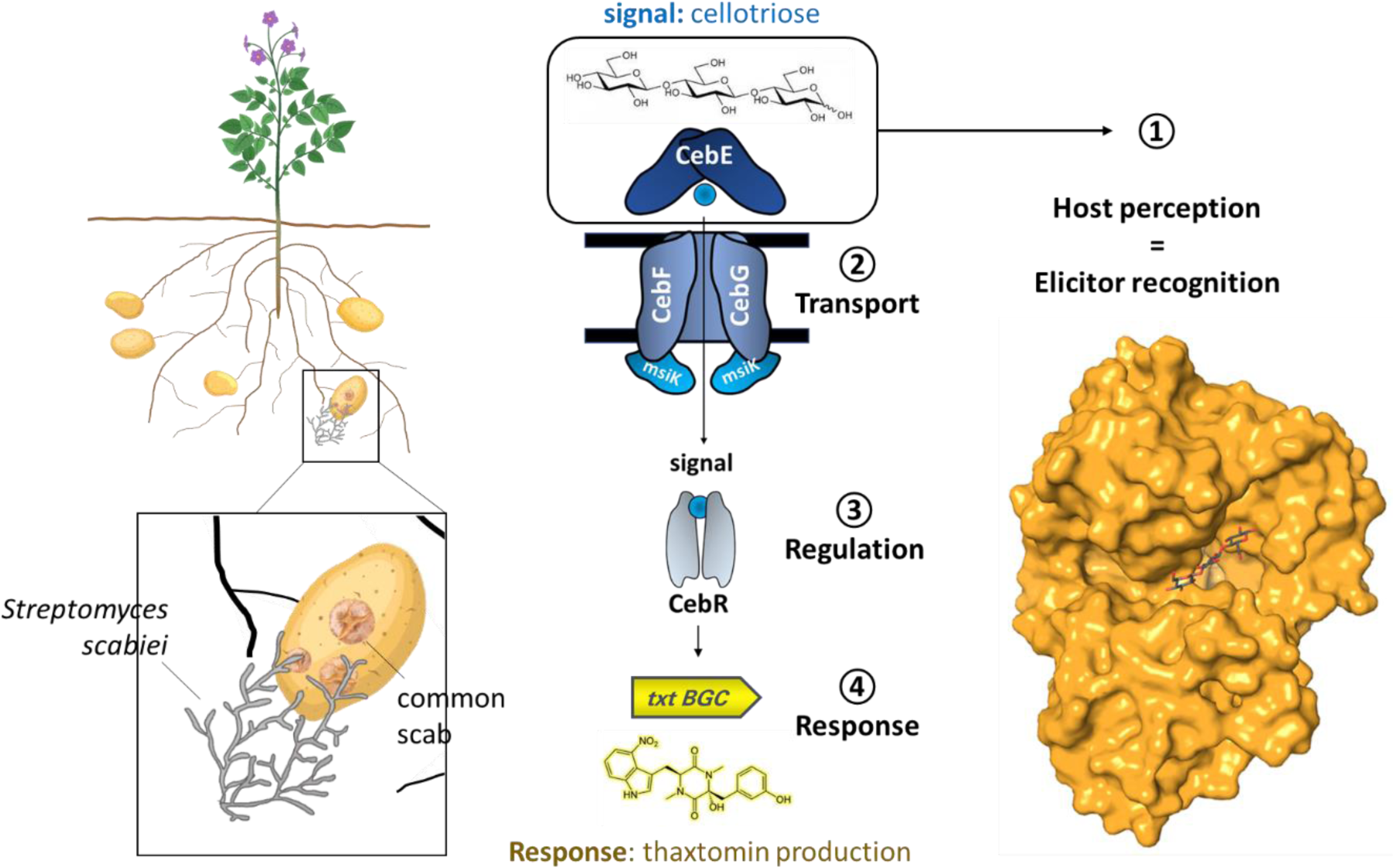

**Highlights:**

- Cellotriose uptake triggers common scab in tuber/root crops by *Streptomyces scabiei*
- Crystal structure of CebE of *S*. *scabiei* interacting with cellotriose is solved
- Cellotriose triggers the closed conformational state of CebE
- The CebE/cellotriose route to pathogenicity is conserved in *Streptomyces* species
- CebE-type background may affect the cellotriose concentration eliciting virulence

## 1. Introduction

Microbes interacting with plants, either beneficial or pathogenic, must perceive molecules indicating the presence of their host to trigger the appropriate response for tissue penetration and colonization (1). Protein-ligand interactions are also often the starting event for saprophytic microorganisms who must sense byproducts released from the decaying plant material to switch on the expression of specific carbon source uptake and catabolic systems. For organic soil-dwelling bacteria, lignocellulose is a major nutrient reservoir, first by itself for being the most abundant polysaccharide on earth, but also because crystalline cellulose is physically linked to other important nourishing polymers such as xylan, mannan, pectin, and lignin. Most members of the bacterial genus *Streptomyces* have acquired a complete cellulolytic system that comprises structurally diverse and synergistically acting secreted cellulose degrading enzymes to generate, import, and consume cellobiose and other cello-oligosaccharides (2–5). In these species, cellobiose - the main carbohydrate released by the cellulolytic system (6–8) - and cellotriose, are actively imported by the CebEFG-MsiK ATP-binding cassette (ABC) transporter (9–12). CebE is the sugar-binding component of the ABC transporter, proteins CebF and CebG form the transporter permease, and the energy for active cello-oligosaccharide transport is provided by the multiple sugar importer ATPase MsiK. The imported cello-oligosaccharides are subsequently hydrolyzed by the main beta-glucosidase BglC, and/or the alternative beta-glucosidase BcpE1 (13) to feed the glycolysis directly with glucose (14). The use of carbohydrates emanating from cellulose degradation appears to be so crucial for catabolism that multiple copies of the *cebR*-*cebEFG*-*bglC* gene cluster are often present in the genomes of these organisms, either acquired by horizontal transfer (xenologs) or by gene duplication (paralogs) (3, 10, 13, 15–17).

For *Streptomyces scabiei* (syn. *scabies*) and other *Streptomyces* species producing the thaxtomin phytotoxins responsible for the disease called common scab on root and tuber crops, cello-oligosaccharides emanating from the plant cell wall are not only perceived as nutrients but also as signals for triggering their pathogenic lifestyle (9, 15, 18–20). Indeed, the production of thaxtomins and other key metabolites of the virulome is activated by the transport of cello-oligosaccharides, particularly cellotriose (15, 18, 21, 22). The binding-affinity of CebE of *S*. *scabiei* (CebE^scab^) for cello-oligosaccharides has *K*_D_ values of 14 (±2) nM and 2 (±0.5) nM for cellobiose and cellotriose, respectively (9). Instead, the affinity for cello-oligosaccharides of the CebE protein of the highly cellulolytic species *Streptomyces reticuli* (CebE^reti^) is much lower with *K*_D_ at the micromolar level (23). The high affinity of CebE^scab^ to cello-oligosaccharides would make *S*. *scabiei* one of the first beneficiaries of the cello-oligosaccharides released by efficient lignocellulolytic microorganisms. Yet, for this strain that evolved to a disabled cellulolytic system (18, 24) it would be crucial to be able to distinguish between cello-oligosaccharides from living and decaying plant material. We postulated that the particularly high affinity of CebE^scab^ for cellotriose could be a key feature of *S*. *scabiei* for discerning living plants from plant decaying material and therefore to adopt either a pathogenic or saprophytic lifestyle (24). Indeed, cellobiose is by far the main product released by the cellulolytic systems (6–8), while Johnson et al. detected only cellotriose released from rapidly growing radish seedling and from actively dividing tobacco NT1 cells in suspension (18). Therefore, cellotriose is thought to be perceived as the signal molecule specifying *S*. *scabiei* the nearby presence of a growing host to colonize, whereas sensing cellobiose would indicate *S*. *scabiei* dead plant material to consume (15, 24). In addition, *S*. *scabiei* species possess the alternative CebEFG2 ABC-transporter system that also participates in the uptake of cello-oligosaccharide elicitors (17).

In this work, we elucidated the structure of CebE^scab^ in complex with cellotriose thereby identifying the key residues involved in elicitor recognition for the onset of the pathogenic lifestyle of *S*. *scabiei* and other phytopathogenic *Streptomyces* species.

## 2. Materials and methods

### 2.1. Strains, chemicals, and culture conditions

*Escherichia coli* DH5α was used for routine molecular biology applications, and *Escherichia coli* BL21(DE3) Rosetta™ (Novagen) for heterologous protein production. *E*. *coli* strains were cultured in LB (BD Difco LB broth) medium supplemented with the appropriate antibiotics (kanamycin [50 μg/ml], chloramphenicol [25 μg/ml]). Cellotriose was purchased from Carbosynth.

### 2.2. Production and purification of CebE from *S*. *scabiei* 87-22

Heterologous production of CebE^scab^ (SCAB_57751; WP_013003368) was performed in strain *E*. *coli* BL21(DE3) Rosetta™ (Novagen) harboring pSAJ016 (pET28a derivative containing the coding sequence of *scab57751* (*cebE*) without the first 132 nt – corresponding to the signal peptide – inserted into NdeI and HindIII restriction sites (9)). Production and purification by nickel affinity chromatography were performed as previously described (9).

#### Crystallization and structure determination of CebE^scab^ in complex with cellotriose

CebE^scab^ was concentrated to 15 mg/ml in a Tris-HCl 30 mM pH 7.5 buffer containing 150 mM NaCl. Cellotriose was added to a 15 mM final concentration. Crystals were obtained using the sitting-drop vapor diffusion method at 4°C with drops made of 0.2 μl of protein solution mixed with 0.2 μl of precipitant solution (polyethylene glycol 3350 25% w/v, and sodium citrate buffer 0.1 M pH 3.5). The crystal was transferred into a cryoprotectant solution containing 45% v/v glycerol and 20% w/v polyethylene glycol 6000 before flash-freezing in a liquid nitrogen bath. Diffraction data were collected at the Soleil Synchrotron PROXIMA 2A beamline (Paris). Data were integrated and scaled using XDS (25). Initial phases were obtained by molecular replacement with the AlphaFold (26) model of CebE^scab^ as search model using Phaser (27). The structure was built with Coot (28) and refined with Refmac (29). The figures were prepared using PyMOL (The PyMOL Molecular Graphics System, Version 2.4.1 Enhanced for Mac OS X, Schrödinger, LLC.). The CebE structure in complex with cellotriose can be found at PDB DOI:https://www.rcsb.org/structure/8BFY.

#### Regulon predictions

Computational prediction of CebR binding sites was performed with the PREDetector software (30) according to the methodology and philosophy described in (31). Sequences used to generate the CebR position weight matrix are available at https://github.com/SebaRigali/PWMs/blob/main/CebR%20PWM%20230412.

## Results and discussion

### Overall three-dimensional structure of CebE^scab^ binding cellotriose

We obtained the crystallographic structure of CebE^scab^ in complex with cellotriose at a 1.55 Å resolution. The crystal belongs to the P2_1_ space group with one molecule in the asymmetric unit. The final R_work_ and R_free_ are 13.6% and 17.8%, respectively (Table 1). The CebE^scab^ structure contains residues 62 to 454. A single segment of 33 amino acids at the N-terminus, which includes 17 residues from the CebE sequence and 16 residues from the His-Tag used for purification, has not been modeled because of lack of electron density.

**Table 1.**
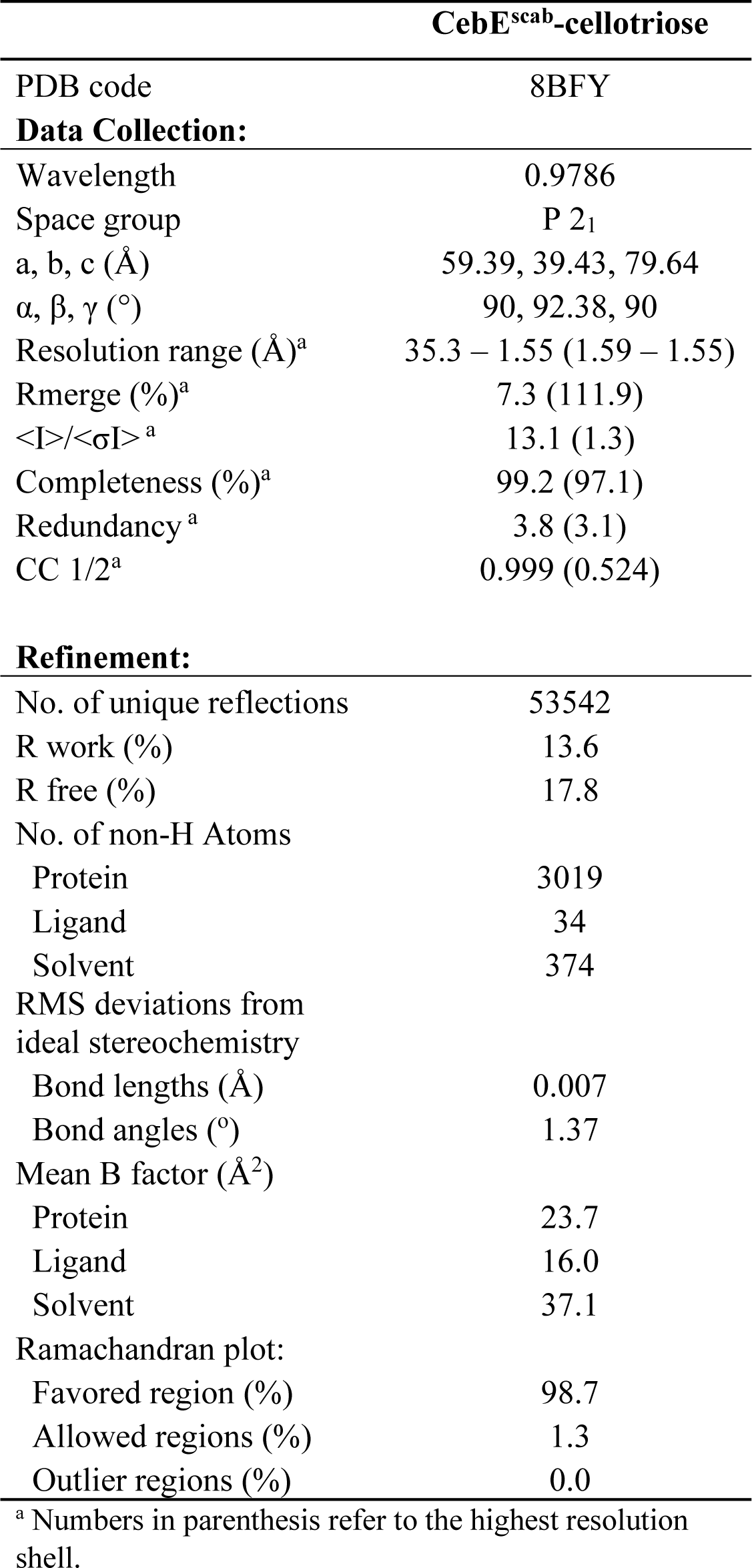
Data collection and refinement statistics.

| **CebE<sup>scab</sup>-cellotriose** |  |
| --- | --- |
| PDB code | 8BFY |
| **Data Collection:** |  |
| Wavelength | 0.9786 |
| Space group | P 2<sub>1</sub> |
| a, b, c (Å) | 59.39, 39.43, 79.64 |
| $\alpha$, $\beta$, $\gamma$ (°) | 90, 92.38, 90 |
| Resolution range (Å)<sup>a</sup> | 35.3 – 1.55 (1.59 – 1.55) |
| Rmerge (%)<sup>a</sup> | 7.3 (111.9) |
| $\langle I \rangle / \langle \sigma I \rangle$<sup>a</sup> | 13.1 (1.3) |
| Completeness (%)<sup>a</sup> | 99.2 (97.1) |
| Redundancy<sup>a</sup> | 3.8 (3.1) |
| CC 1/2<sup>a</sup> | 0.999 (0.524) |
| **Refinement:** |  |
| No. of unique reflections | 53542 |
| R work (%) | 13.6 |
| R free (%) | 17.8 |
| No. of non-H Atoms |  |
| Protein | 3019 |
| Ligand | 34 |
| Solvent | 374 |
| RMS deviations from ideal stereochemistry |  |
| Bond lengths (Å) | 0.007 |
| Bond angles (°) | 1.37 |
| Mean B factor (Å<sup>2</sup>) |  |
| Protein | 23.7 |
| Ligand | 16.0 |
| Solvent | 37.1 |
| Ramachandran plot: |  |
| Favored region (%) | 98.7 |
| Allowed regions (%) | 1.3 |
| Outlier regions (%) | 0.0 |

<sup>a</sup> Numbers in parenthesis refer to the highest resolution shell.

CebE^scab^ adopts a cluster B type SBP-fold as described by Bemtsson et al. (32, 33). It is composed of two domains connected by three hinge regions. Domain 1 contains residues 62-176 and 330-386, and is made of a six-stranded β-sheet surrounded by ten helices (Figure 1). The larger Domain 2 includes residues 177-329 and 387-454, and is formed by a four-stranded β-sheet surrounded by ten helices with the two C-terminal helices packed on it. The elongated ligand binding pocket is located at the interface between the two domains (Figure 1B).

**Figure 1.**
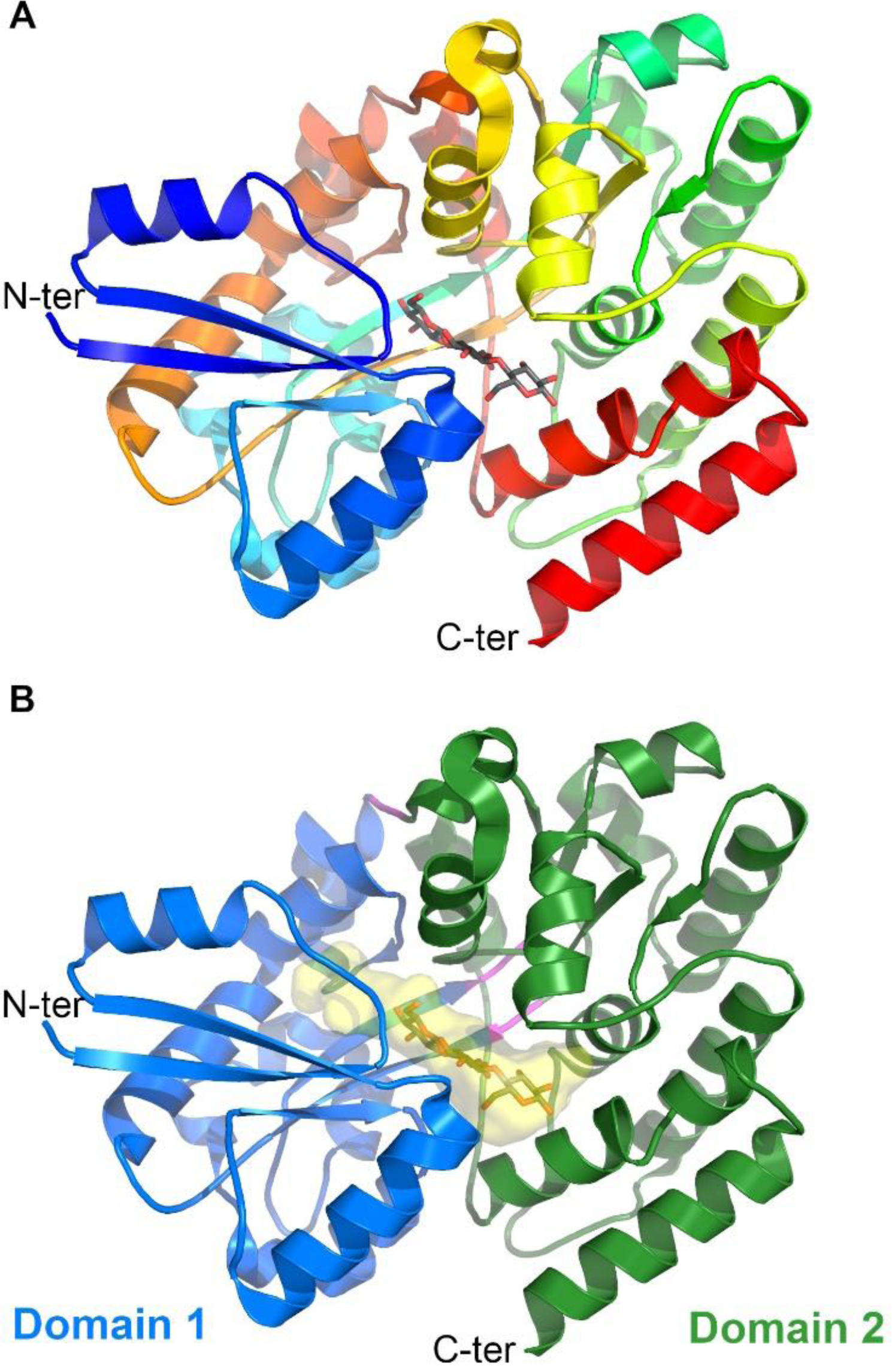
Overall structure of the CebE/cellotriose complex. **A**. Cartoon representation of CebE^scab^ with a rainbow coloring from the N-terminus in blue to the C-terminus in red. The cellotriose molecule is displayed as gray sticks. **B.** Cartoon representation of CebE^scab^ with Domains 1 and 2 in blue and green, respectively. The three hinge regions are colored in magenta. The closed cavity occupied by cellotriose (gray sticks) is shown as a yellow transparent surface.

### The cellotriose binding site of CebE^scab^

CebE^scab^ was crystalized in the presence of 15 mM cellotriose, which approximately corresponds to a 50-fold excess compared to the protein concentration. An electron density corresponding to the whole cellotriose molecule is observed in the ligand binding pocket (Figure 2A), clearly establishing the two β-1,4 links between the three β-D-glucoses. The reducing end (D-Glc1) is more inserted in Domain 2 whereas the non-reducing end (D-Glc3) makes more interactions with Domain 1 (Table 2). Cellotriose is stabilized in the pocket by few hydrophobic interactions and numerous H-bonds, nine of them being mediated by eight water molecules surrounding the ligand (Figure 2B and C, Table 2). D-Glc3 provides the highest contribution to the binding of cellotriose, being involved in ten H-bonds and three hydrophobic interactions with the sidechain of W303 (parallel stacking), F70, and M123. D-Glc1 and D-Glc2 follow with 7 and 6 H-bonds, respectively, the latter being involved in an additional hydrophobic interaction with F282.

**Figure 2.**
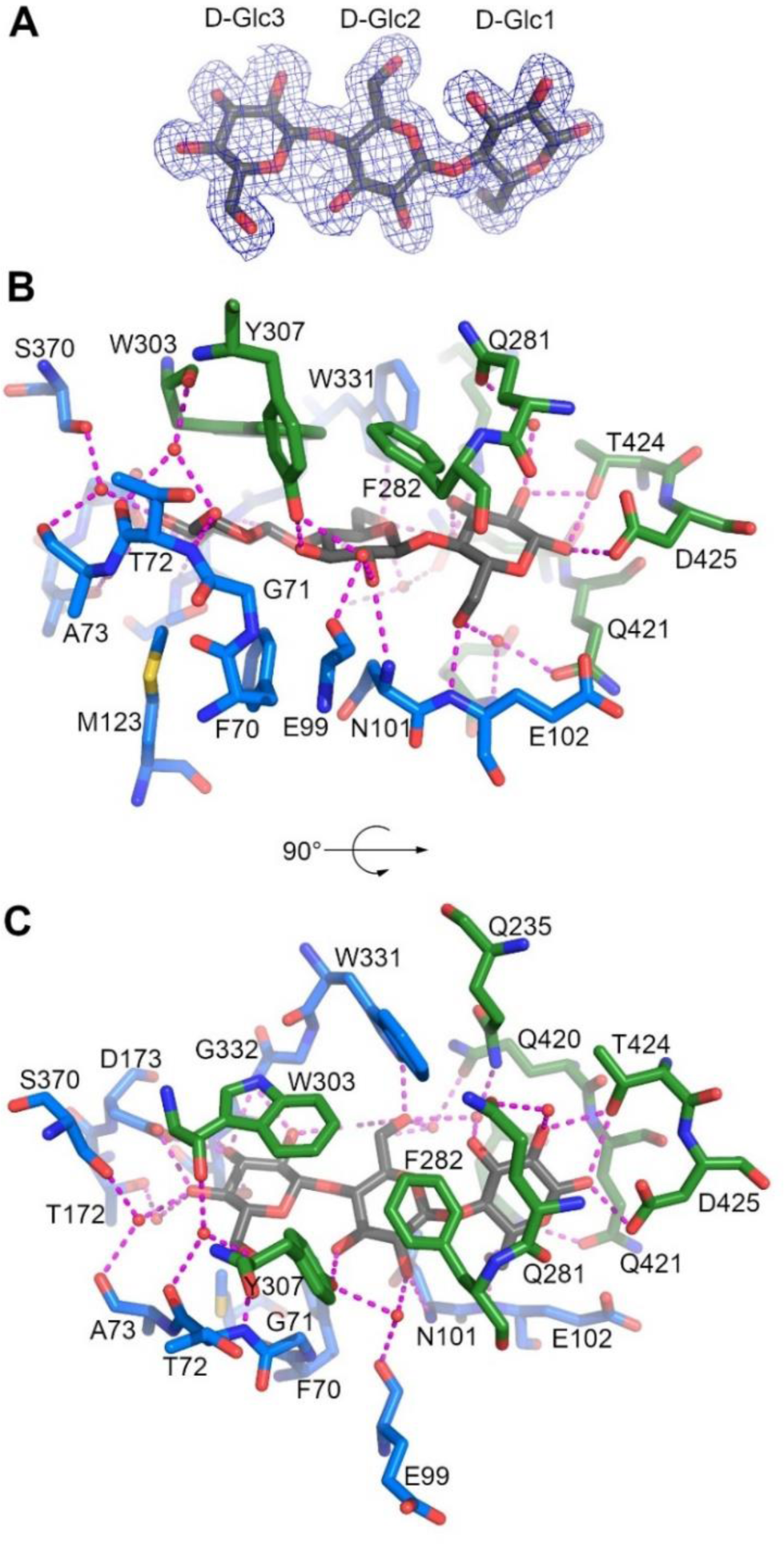
Substrate binding site of the CebE:cellotriose complex. **A**. 2Fo-Fc electron density map displayed at 1 σ level around cellotriose. **B**. Interactions stabilizing cellotriose (gray sticks), residues from Domains 1 and 2 are displayed as blue and green sticks respectively, water molecules as small red spheres, and H-bonds as magenta dashed lines. **C**. same as **B** with a 90° rotation around a horizontal axis.

**Table 2.** List of interactions between CebE^scab^ and cellotriose.

| Cellotriose monomer | Cellotriose atom | CebE domain | CebE residue | CebE atom | Distance (Å) | Type of interaction |
| --- | --- | --- | --- | --- | --- | --- |
| D-Glc1 | O1 | 2 | D425 | OD2 | 2.8 | H-bond |
|  | O1 | 2 | T424 | OG1 | 3.1 | H-bond |
|  | O2 | 2 | T424 | OG1 | 2.8 | H-bond |
|  | O2 | 2 | Q281 | OE1 | 2.8/2.7 | H<sub>2</sub>O mediated H-bond* |
|  | O3 | 2 | Q235 | NE2 | 2.8/3.0 | H<sub>2</sub>O mediated H-bond* |
|  | O6 | 1 | E102 | N | 2.8 | H-bond |
|  | O6 | 2 | Q417 (Q421) | NE2 (OE1) | 2.7/2.9 (2.6) | H<sub>2</sub>O mediated H-bond* |
| D-Glc2 | O2 | 1 | N101 | N | 3.2 | H-bond |
|  | O2 | 1 | E99 (Y307) | O (OH) | 2.7/2.7 (2.9) | H<sub>2</sub>O mediated H-bond* |
|  | O3 | 2 | Y307 | OH | 2.7 | H-bond |
|  | O6 | 1 | W331 | NE2 | 2.9 | H-bond |
|  | O6 | 2 | Q420 (E125) | OE1 (OE1) | 3.1/2.7 (2.6) | H<sub>2</sub>O mediated H-bond* |
|  | O6 | 2 | Q235 | NE2 | 2.9/3.0 | H<sub>2</sub>O mediated H-bond* |
|  | ring | 2 | F282 | phenyl | 4.7 | hydrophobic |
| D-Glc3 | O2 | 1 | E125 | OE2 | 2.7 | H-bond |
|  | O2 | 1 | G333 | N | 2.9 | H-bond |
|  | O3 | 1 | G333 | N | 3.0 | H-bond |
|  | O3 | 1 | D173 | OD1 | 2.6 | H-bond |
|  | O3 | 1 | S334 | OG | 2.7 | H-bond |
|  | O4 | 1 | D173 | OD2 | 2.7 | H-bond |
|  | O4 | 1 | T172 | OG1 | 3.0/2.8 | H<sub>2</sub>O mediated H-bond* |
|  | O4 | 1 | S370<br>(A73) | OG (O) | 2.8/2.5<br>(2.9) | H<sub>2</sub>O mediated H-bond* |
|  | O6 | 1 | T72 | N | 2.8 | H-bond |
|  | O6 | 2 | W303<br>(T72) | O (O) | 2.7/2.7<br>(2.9) | H<sub>2</sub>O mediated H-bond* |
|  | ring | 2 | W303 | indole | 4.1 | hydrophobic |
|  | ring | 1 | F70 | phenyl | 3.7 | hydrophobic |
|  | ring | 1 | M123 | CE | 4.8 | hydrophobic |

\* When H<sub>2</sub>O mediated H-bond, the first distance is related to the cellotriose-H<sub>2</sub>O bound and the second to the H<sub>2</sub>O-protein bound. Parentheses are used when a second residue of the protein contributed to the binding of the H<sub>2</sub>O molecule. Lines shaded in grey highlight the residues of CebE<sup>scab</sup> that are substituted in CebE<sup>reti</sup>.

The importance of D-Glc3 for the affinity of cellotriose is further highlighted by comparing the CebE^scab^:cellotriose structure with the closest structure of SBP proteins in complex with a ligand available in the Protein Data Bank (Figure S1): the ABC transporter associated binding protein from *Bifidobacterium animalis* (Bal6GBP) in complex with β-1,6-galactobiose (PDB code 6H0H, 26% of sequence identity (34)), the ABC transporter associated binding protein AbnE from *Geobacillus stearothermophilus* in complex with arabinohexaose (PDB code 6RKH, 26% of sequence identity (35)) and the galacto-N-biose-/lacto-N-biose I-binding protein (GL-BP) of the ABC transporter from *Bifidobacterium longum* in complex with lacto-N-tetraose (36). Indeed, despite the difference in length and composition of the different oligosaccharides present in the three structures, a saccharide is always bound at a position equivalent to that of D-Glc3, and two residues important for its binding, D173 and W303, are conserved in the structure of these four different ABC-type sugar-binding proteins (Figure S1), as well as in CebE2, the alternative cello-oligosaccharide transporter of *S*. *scabiei* strains (17). While there seems to be a preference for the nonreducing end of the oligosaccharide, it is not exclusive as illustrated by the AbnE:arabinohexaose complex in which it is the fifth arabinose that is bound at this conserved binding position.

### Substrate induced closing of the CebE^scab^ pocket

In the CebE^scab^:cellotriose structure, only seven H-bonds are observed between residues of Domains 1 and 2 when the three hinge regions are removed (P176-M177, G329-N330 and A386-K387), involving residues R100, G127, N128, E131, W331 and Q368 for Domain 1, and Q281, F282, W303, K309, K415 and Q417 for Domain 2. In addition, the two significant hydrophobic interactions between the domains are located in the vicinity of the three hinge regions (P302, F400 and I396 with F365 and F371 for the first hydrophobic cluster, and V409 and I407 with W155 for the second). It therefore seems that an important part of the interactions between Domains 1 and 2 are mediated by the cellotriose molecule itself. The ligand would therefore be responsible for the closing of the sugar-binding pocket, a phenomenon termed as induced-fit ligand binding mechanism (37). This is compatible with the known flexibility of these domains around the hinge regions (35), which is necessary for the cavity to open and allow access to the ligand. Indeed, in the CebE^scab^:cellotriose structure, the pocket accommodating the cellotriose molecule has no access to the solvent (Figure 1B).

A good estimate of the magnitude of the opening can be obtained by comparing our closed CebE^scab^ structure with the structure of the Solute Binding Domain protein from *Kribbella flavida* DSM 17836 (KfSBP, PDB code 5IXP) (Figure 3). KfSBP is the closest homolog of CebE^scab^ in the Protein Data Bank and shares 38% of sequence identity with CebE^scab^, including most of the interactions with cellotriose. Nine residues with sidechain interacting with the ligand differ between KfSBP and CebE^scab^: the M123A mutation is compensated by the A73F one, the F282W substitution provides a more extended hydrophobic interaction but it is conjugated with the Y307V mutation that induces the loss of a H-bond with cellotriose, the D425E difference should maintain the H-bond, and the Q235E, Q281A, Q417G, and Q420N substitutions induce modifications of the water mediated H-bond network that are difficult to quantify. When the two Domain 1 of the two proteins are superimposed, the two Domain 2 are separated by a rotation around the hinge regions of approximately 40° (Figure 3), inducing a large opening of the ligand binding pocket. The extent of the conformational change is better perceived with the Supplemental movie 1 displaying a morphing between the CebE^scab^:cellobiose structure and a CebE^scab^ model obtained using the comparative modeling method RosettaCM (38) with the KfSBP structure as template.

**Figure 3.**
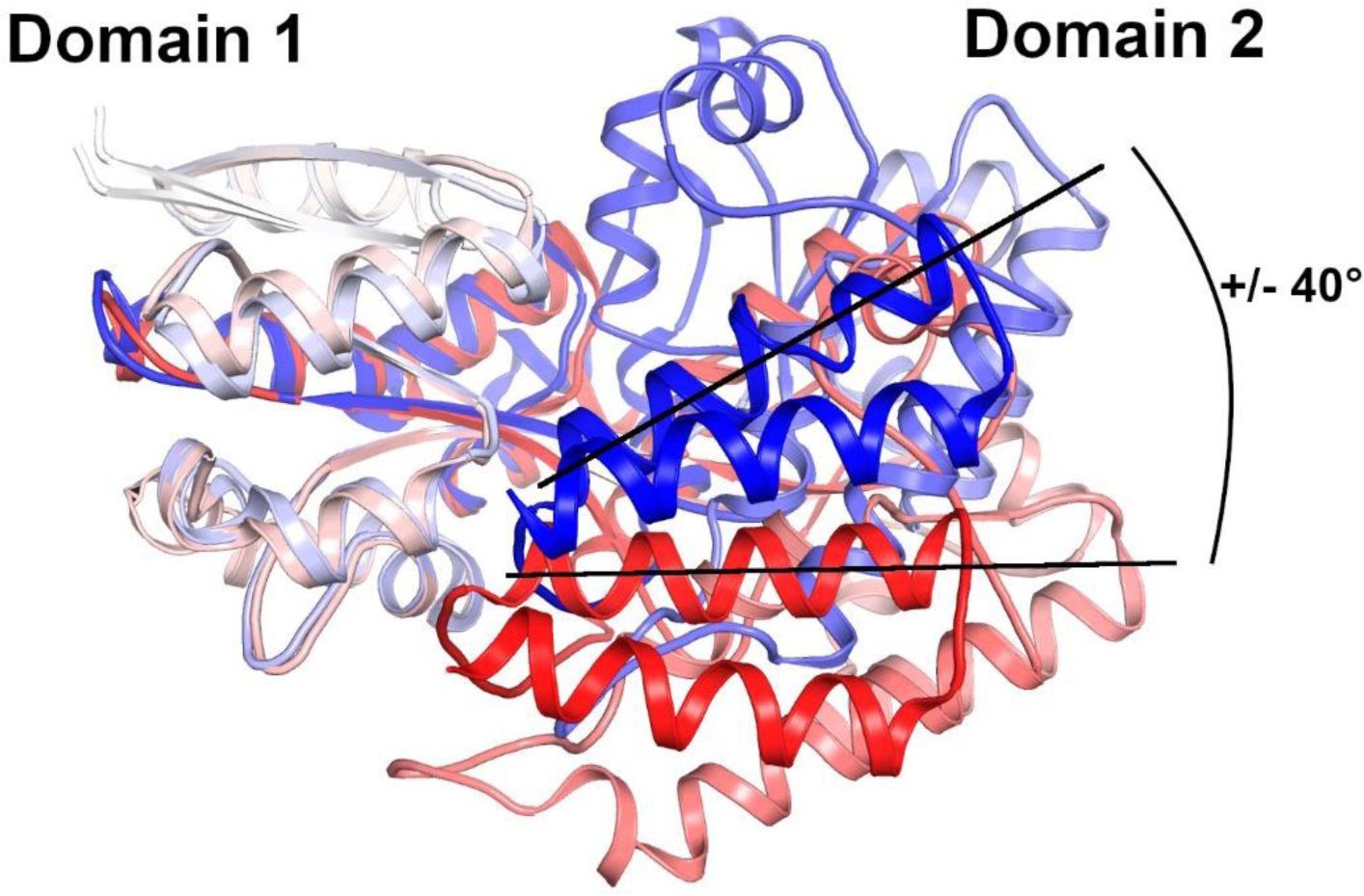
Conformational change of CebE upon cellotriose binding. Superimposition of the CebE^scab^:cellotriose structure (colored with a gradient from the white N-terminus to the blue C-terminus) to the KfSBP structure in an open conformation (colored with a gradient from the white N-terminus to the red C-terminus); only Domain 1 residues were used to calculate the superimposition. Note the +/- 40° angle corresponding to the movement of Domain 2 upon cellotriose-binding.

### Conservation of the cellotriose-mediated signaling pathway in pathogenic *Streptomyces* species

The ABC transport systems for cello-oligosaccharide import by both pathogenic and saprophytic streptomycete species are clustered in at least four different paralog/xenolog subgroups (17), namely CebE^scab^ (9), CebE2^scab^ (17), CebE^reti^ (10, 23), and CebE^gris^ (16). CebE^scab^ and CebE^reti^ are able to bind cellotriose at nano- and micro-molar ranges, respectively, while the *K*_D_ values of CebE^gris^ and CebE2^scab^ for cellobiose and cello-oligosaccharides have not been experimentally determined (9, 10). The genes required for producing thaxtomin phytotoxins, which are included in the pathogenicity island that has been horizontally transferred to different saprophytic *Streptomyces* species, have therefore been integrated into genomes with different backgrounds regarding the affinity of the CebE protein for its substrates. Table 3 lists all thaxtomin-producing pathogenic *Streptomyces* species (57 strains from 10 different species) for which a good quality genome sequence was available and therefore where we could identify the type(s) of CebE protein(s) involved in cellotriose and cellobiose uptake. In addition, the assessment of the conservation of the cellotriose-mediated induction of pathogenicity in all selected strains, was performed by screening for binding sites of the transcriptional repressor for cellulose and cello-oligosaccharide utilization CebR in the biosynthetic gene cluster associated with thaxtomin production (*txt* cluster), and within the *cebEFG* operon. Two main groups can be distinguished, i.e., the species that possess CebE^scab^ (44 strains from 8 different species), and those that possess CebE^reti^ (13 strains from 2 species). Surprisingly, none of the pathogenic strains (with genome available) recruited the CebE-like protein of *Streptomyces griseus* group (16) as elicitor importer. The CebE^scab^ group includes strains that belong to species *S. scabiei*, *S. acidiscabies*, *S. europeiscabiei*, *S. stelliscabiei*, *S. brasiliscabiei*, *S. griseiscabiei*, and *S. niveiscabiei* (Table 3). *S*. *ipomoeae* is also part of this group, but the absence of a CebR-binding site within the *txt* cluster would explain earlier results that suggested this species did not select the CebE-cello-oligosaccharide-mediated pathway for induction of thaxtomin production (39, 40). The CebE^reti^ group includes the strains that belong to species *S. turgidiscabies*, and *S. caniscabiei* (Table 3).

**Table 3.** Type of cellotriose/CebE-mediated signaling pathways to thaxtomin production.

| Pathogenic *Streptomyces* |  | Thaxtomin biosynthesis |  |  | CebE subgroup |  |  | CebR-binding sites<sup>3</sup> |  |  |  |
| --- | --- | --- | --- | --- | --- | --- | --- | --- | --- | --- | --- |
| species | strain | TxtA<sup>1</sup> | TxtB<sup>1</sup> | TxtR<sup>1</sup> | *scabiei*<sup>2</sup> | *reticuli*<sup>2</sup> | *griseus*<sup>2</sup> | *txtR* | *txtA* | *txtB* | *cebEFG* |
| *scabiei* | 87-22 | 100 | 100 | 100 | 100 | 45 | 41 | 20.8 | 20.8 | 20 | 23 |
| *scabiei* | 84-34 | 98 | 100 | 100 | 100 | 45 | 41 | 21 | 21 | 20.2 | 23 |
| *scabiei* | 84-232 | 100 | 100 | 100 | 100 | 45 | 41 | 21 | 21 | 20.3 | 23 |
| *scabiei* | 85-08 | 100 | 100 | 100 | 100 | 45 | 41 | 21 | 21 | 20.2 | 23 |
| *scabiei* | 95-18 | 100 | 100 | 100 | 100 | 45 | 41 | 21 | 21 | 20.3 | 23 |
| *scabiei* | 96-06 | 98 | 100 | 100 | 100 | 45 | 41 | 21 | 21 | 20.3 | 23 |
| *scabiei* | 96-15 | 100 | 100 | 100 | 100 | 45 | 41 | 21 | 21 | 20.2 | 23 |
| *scabiei* | LBUM 1475 | 100 | 100 | 100 | 100 | 45 | 41 | 20.8 | 12.6 | 20 | 23 |
| *scabiei* | LBUM 1477 | 99 | 100 | 100 | 100 | 45 | 41 | 20.8 | 20.8 | 20 | 23 |
| *scabiei* | LBUM 1478 | 100 | 100 | 100 | 100 | 45 | 41 | 20.8 | 20.8 | 20.1 | 23 |
| *scabiei* | LBUM 1479 | 99 | 97 | 100 | 100 | 45 | 41 | 20.8 | 20.8 | 20 | 23 |
| *scabiei* | LBUM 1480 | 100 | 100 | 100 | 100 | 45 | 41 | 20.8 | 20.8 | 20 | 23 |
| *scabiei* | LBUM 1481 | 99 | 100 | 100 | 100 | 45 | 41 | 20.8 | 20.8 | 20 | 23 |
| *scabiei* | LBUM 1482 | 100 | 100 | 100 | 100 | 45 | 41 | 20.8 | 20.8 | 20 | 23 |
| *scabiei* | LBUM 1483 | 99 | 100 | 100 | 100 | 45 | 41 | 20.8 | 20.8 | 20 | 23 |
| *scabiei* | LBUM 1484 | 98 | 100 | 100 | 100 | 45 | 41 | 20.8 | 20.8 | 20 | 23 |
| *scabiei* | LBUM 1485 | 98 | 100 | 100 | 100 | 45 | 41 | 20.8 | 20.8 | 20 | 23 |
| *scabiei* | LBUM 1486 | 99 | 100 | 100 | 100 | 45 | 41 | 20.8 | 20.8 | 20 | 23 |
| *scabiei* | LBUM 1487 | 99 | 100 | 100 | 100 | 45 | 41 | 20.8 | 20.8 | 20 | 23 |
| *scabiei* | LBUM 1488 | 99 | 100 | 100 | 100 | 45 | 41 | 20.8 | 20.8 | 20 | 23 |
| *scabiei* | NCPPB 4066 | 98 | 99 | 100 | 100 | 45 | 41 | 20.8 | 20.8 | 20 | 23 |
| *scabiei* | NRRL B-16523 | 98 | 100 | 100 | 100 | 45 | 41 | 21 | 21 | 20.3 | 23 |
| *scabiei* | St129 | 99 | 100 | 100 | 100 | 45 | 41 | 21.1 | 12.7 | 20.1 | 23 |
| *brasiliscabiei* | IBSBF2867 | 99 | 100 | 100 | 89 | 46 | 41 | 20.9 | 20.9 | 20.2 | 23 |
| *griseiscabiei* | NRRL B-2795 | 100 | 100 | 100 | 84 | 46 | 41 | 20.8 | 20.8 | 20 | 23 |
| *ipomoeae* | 91-03 | 49 | 61 | 42 | 80 | 46 | 44 | 8.3 | 8.3 | 5.8 | 23 |
| *ipomoeae* | 78-51 | 49 | 61 | 51 | 80 | 45 | 44 | 8.2 | 8.6 | 5.8 | 23 |
| *ipomoeae* | 88-35 | 49 | 61 | 42 | 72 | 45 | 44 | 8.2 | 8.6 | 5.9 | 23 |
| *scabiei* | NCPPB 4086 | 100 | 100 | 97 | 72 | 45 | 44 | 20.8 | 20.8 | 16.7 | 23 |
| *europaeiscabiei* | 96-14 | 99 | 100 | 99 | 72 | 45 | 44 | 20.9 | 20.9 | 20.2 | 23 |
| *europaeiscabiei* | St1229 | 99 | 100 | 99 | 72 | 45 | 44 | 20.9 | 20.9 | 20.2 | 23 |
| *europaeiscabiei* | NRRL B-24443 | 98 | 100 | 100 | 72 | 45 | 44 | 21 | 21 | 20.2 | 23 |
| *europaeiscabiei* | 89-04 | 98 | 100 | 100 | 72 | 45 | 44 | 20.9 | 20.9 | 20.2 | 23 |
| *stelliscabiei* | DSM 41803 | 99 | 100 | 98 | 70 | 45 | 43 | 20.8 | 20.8 | 20.1 | 23 |
| *stelliscabiei* | P3825 | 98 | 100 | 99 | 70 | 45 | 43 | 20.8 | 20.8 | 20.1 | 23 |
| *stelliscabiei* | NRRL B-24447 | 99 | 100 | 99 | 70 | 45 | 43 | 21 | 21 | 20.3 | 23 |
| *acidiscabies* | 84-104 | 100 | 100 | 100 | 68 | 51 | 42 | 20.9 | 12.7 | 20.1 | 23 |
| *acidiscabies* | a10 | 100 | 100 | 100 | 68 | 51 | 42 | 20.9 | 20.9 | 20.2 | 23 |
| *acidiscabies* | 98-49 | 100 | 100 | 100 | 68 | 51 | 42 | 20.9 | 20.9 | 20.2 | 23 |
| *acidiscabies* | 85-06 | 100 | 100 | 100 | 68 | 51 | 42 | 20.9 | 20.9 | 20.2 | 23 |
| *acidiscabies* | FL01 | 100 | 100 | 99 | 68 | 51 | 42 | 21.1 | 12.7 | 20.1 | 23 |
| *acidiscabies* | St105 | 100 | 100 | 100 | 68 | 51 | 42 | 20.9 | 20.9 | 20.1 | 23 |
| *acidiscabies* | LBUM 1476 | 100 | 100 | 100 | 68 | 51 | 42 | 20.8 | 20.8 | 20.1 | 23 |
| *niveiscabiei* | NRRL B-24457 | 100 | 100 | 100 | 67 | 52 | 43 | 21 | 21 | 20.2 | 23 |
| *turgidiscabies* | T45 | 90 | 90 | 72 | 48 | 80 | 44 | 21.1 | 12.9 | 15.7 | 23 |
| *turgidiscabies* | Car8 | 90 | 90 | 72 | 48 | 75 | 41 | 21 | 12.8 | 15.7 | 23 |
| *caniscabiei* | AMCC400023 | 100 | 100 | 100 | 47 | 79 | 42 | 20.8 | 20.8 | 20 | 23 |
| *caniscabiei* | 96-12 | 99 | 100 | 100 | 47 | 79 | 42 | 21 | 21 | 20.3 | 23 |
| *caniscabiei* | NRRL B-24093 | 99 | 100 | 100 | 47 | 79 | 43 | 21 | 21 | 20.3 | 23 |
| *caniscabiei* | ND05-01C | 99 | 100 | 100 | 47 | 79 | 42 | 20.9 | 20.9 | 20.2 | 23 |
| *caniscabiei* | ND05-13A | 99 | 100 | 100 | 47 | 79 | 42 | 20.9 | 20.9 | 20.2 | 23 |
| *caniscabiei* | ND05-3B | 99 | 100 | 100 | 47 | 79 | 42 | 20.9 | 20.9 | 20.2 | 23 |
| *caniscabiei* | NE06-02D | 99 | 100 | 100 | 47 | 79 | 42 | 20.9 | 20.9 | 20.1 | 23 |
| *caniscabiei* | NE06-02F | 99 | 100 | 100 | 47 | 79 | 43 | 20.9 | 20.9 | 20.2 | 23 |
| *caniscabiei* | ID-03-3A | 99 | 100 | 100 | 47 | 79 | 42 | 20.9 | 20.9 | 20.2 | 23 |
| *caniscabiei* | ID01-6.2a | 99 | 100 | 100 | 47 | 79 | 42 | 20.8 | 20.8 | 20.1 | 23 |
| *caniscabiei* | ID01-12c | 99 | 100 | 100 | 47 | 79 | 42 | 21 | 21 | 20.3 | 23 |

<sup>1</sup>, the values refer to the amino acid identity expressed in % compared to the proteins of *S. scabiei* 87-22 used as reference sequences. <sup>2</sup>, the values refer to the amino acid identity expressed in % compared to the CebE proteins of *S. scabiei* 87-22 (WP\_013003368.1), *S. reticuli* (CAB46342.1), and *S. griseus* (WP\_012379731.1). <sup>3</sup>, the values refer to the score obtained for a 14-nt sequence according to the position weight matrix generated from the experimentally validated CebR-binding sites (23 is the maximum score corresponding to the 14-nt TGGGACGCGTCCCA palindromic sequence).

Based on *K*_D_ values measured for two different CebE protein subgroups, pathogenic *Streptomyces* species would trigger thaxtomin production after sensing cellotriose at either the nano-(CebE^scab^ subgroup) or at the micro-molar level (CebE^reti^ subgroup). This major difference in the CebE affinity for the natural elicitor cellotriose may result in species sensitive to different concentration threshold for the molecules eliciting their pathogenic lifestyle. The molecular origin of this significant difference of affinity can be explored using the AlphaFold (26) model of CebE^reti^ available in the AlphaFold database (UniProt id Q9X9R7_STRRE). This model is of very good quality with an average pLDDT (predicted Local Distance Difference Test) value of 94.3 calculated for the Cα of the globular part of the protein (from I51 to Q444). The CebE^reti^ model corresponds to the closed conformation of the protein and can be very well superimposed to CebE^scab^ (root mean square deviation of 1.4 Å calculate over 387 Ca).

Ten residues of CebE^scab^ directly or indirectly (via a water molecule) involved in twelve interactions with cellotriose are substituted in CebE^reti^ namely, T72V, A73F, E99T, N101T, E102D, M123A, Q235N, Y307Q, Q420N, and Q421T (Table 2, Figure 4). Eight of these ten residues are also substituted (by the same or other amino acids) in the CebE proteins of strains that possess a CebE^reti^ background (Figure 5). Four of these residues, T72, E99, N101, and E102, participate in the H-bond network stabilizing cellotriose through their backbone and their mutation is therefore not expected to affect the binding of the ligand. The loss of hydrophobic interaction resulting from the M123A substitution is compensated by the concomitant A73F substitution. The three glutamines (Q235, Q420, and Q421), located close to each other in the structure, interact with cellotriose via H_2_O-mediated H bonds. Their mutation into asparagine (Q235 and Q420) or threonine (Q421) will slightly modify the H-bond network involving water molecules surrounding the ligand, the effect on binding does however not seem important. The two most significant differences between the CebE^scab^ and CebE^reti^ binding site, potentially explaining the reduced affinity for cellotriose are i) the Y307Q substitution where the loss of the direct H-bond between Y307 and D-Glc2 is due to the substitution by the shorter glutamine residue, and ii) the G127M substitution that brings a hydrophobic side chain in close proximity with a polar area of the ligand, preventing at least one water mediated H-bond. However, the latter two substitutions are only specific to the CebE protein of *S*. *reticuli* and are not conserved in the CebE proteins of the CebE^reti^ subgroup found in the pathogenic species *S. turgidiscabies* and *S. caniscabies* (Figure 5). Therefore, whether pathogenic species with either a CebE^scab^ or CebE^reti^ background would require different concentrations of elicitor for the onset of thaxtomin production will have to be determined experimentally. However, this hypothesis would rather imply the highly dynamic ligand binding mechanism of CebE than residue substitutions specifically involved in cellotriose-binding.

**Figure 4.**
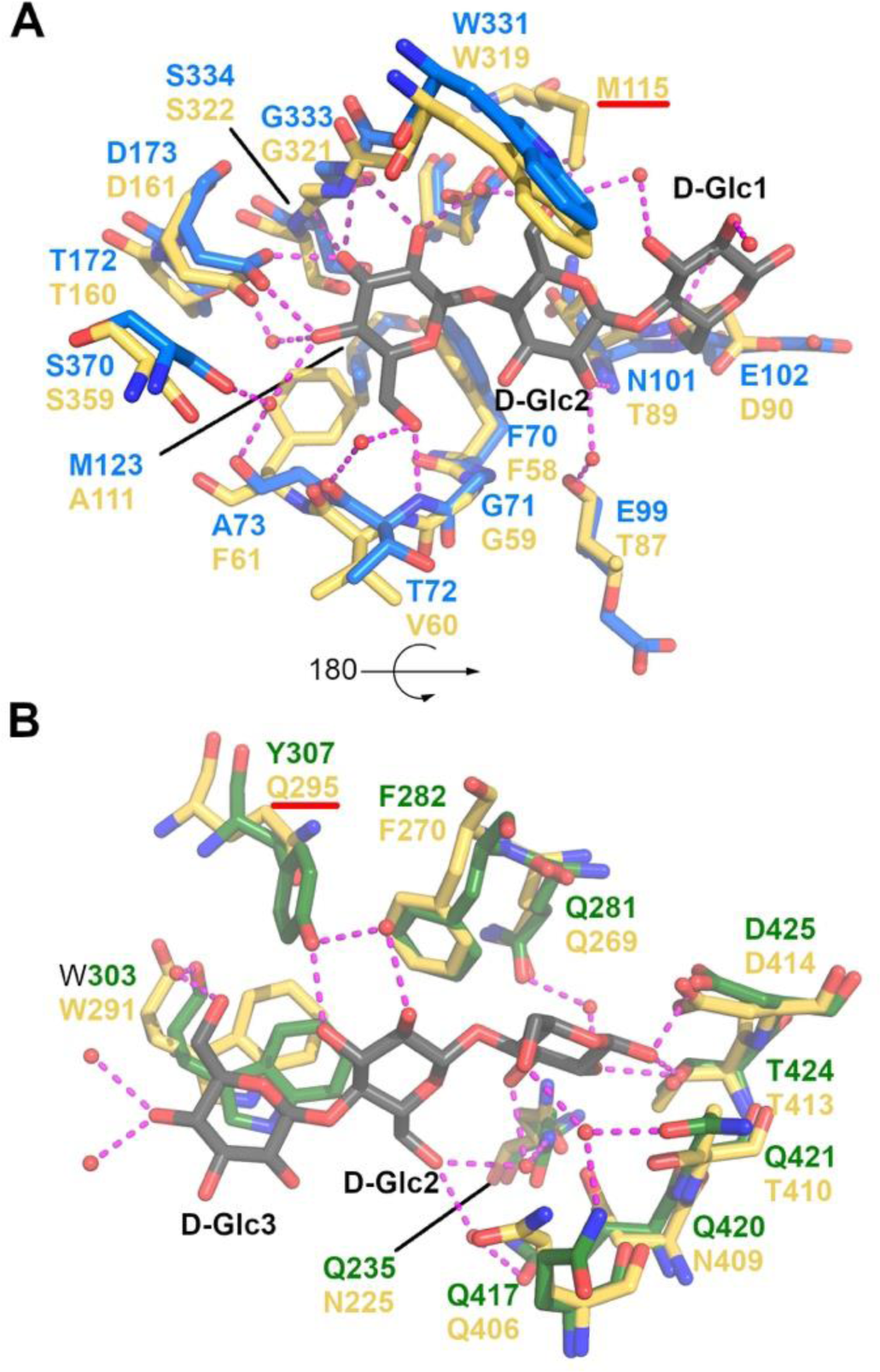
Substrate binding site of the CebE:cellotriose complex superimposed to the CebE^reti^ model. **A**. Interactions of residues from Domain 1 (blue sticks) stabilizing cellotriose (gray sticks) superimposed to their equivalent in CebE^reti^ (yellow sticks), water molecules are displayed as small red spheres, and H-bonds as magenta dashed lines. **B**. same as **A** for domain 2 (green sticks) with a rotation of approximately 180° around a horizontal axis.

**Figure 5.**
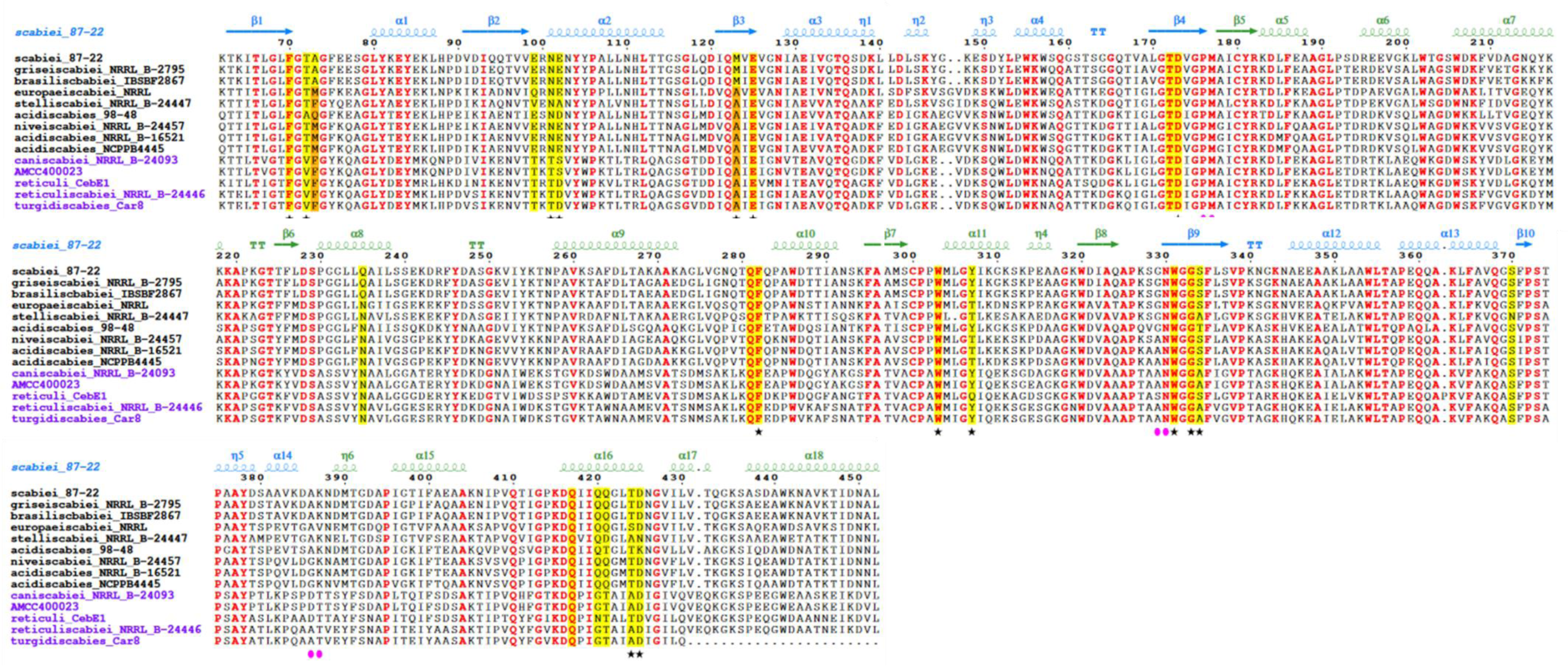
Sequence alignment of CebE proteins from model *Streptomyces* pathogenic species with amino acid numbering of CebE^scab^. Secondary structure elements of the CebE^scab^:cellotriose structure are schematized above the alignment with the same coloring code as in Figure 1B, whereas hinge regions are identified with magenta ellipses below the alignment. Residues strictly conserved are shown in red, and those directly interacting with cellotriose or indirectly interacting with cellotriose via a water molecule are highlighted in yellow (or orange when the short and bulky side chains are switched). Stars indicate direct interaction with the ligand.

## Conclusions

In this work we solved the crystal structure of CebE^scab^ in complex with cellotriose at a resolution of 1.55 Å, thereby revealing the structural basis of the first event responsible for root and tuber plant colonization by *S*. *scabiei*. The interaction between CebE^scab^ and cellotriose involves 26 direct or water-mediated hydrogen bonds and hydrophobic interactions. As previously observed in other sugar-binding proteins of ABC transporters, it is the sugar at the non-reducing end of the oligosaccharide that occupies the most conserved part of the ligand-binding cleft. An induced-fit mechanism is expected to generate the closed conformational changes of CebE, where cellotriose binding triggers the movement between Domains 1 and 2 of the protein. This mechanism is predicted to facilitate the selection between the unliganded and liganded states of SBPs by the transmembrane domains of the importer (37, 41). Prediction of the CebR regulon revealed that the CebE-mediated import of cellotriose is conserved for triggering the production of thaxtomin phytotoxins in pathogenic *Streptomyces* species. The unique loss of the CebR-repressed expression of thaxtomin biosynthetic genes is found in strains belonging to *S*. *ipomoeae* species associated with the colonization of sweet potatoes. Based on sequence similarity between CebE proteins of pathogenic streptomycetes, strains belonging to species *S. acidiscabies*, *S. europeiscabiei*, *S. stelliscabiei*, *S. brasiliscabiei*, *S. S. griseiscabiei*, and *niveiscabiei* would sense the presence of cellotriose with similar affinity as the one previously calculated for CebE^scab^. Instead, pathogenic *Streptomyces* strains of species *S. turgidiscabies* and *S. caniscabies* possess a CebE protein orthologous to CebE^reti^ with lower affinity for cellotriose, suggesting that they could possibly need a higher quantity of cellotriose released by their host to induce the colonization process. However, this hypothesis would imply the highly dynamic ligand binding mechanism of CebE rather than residues specifically involved in cellotriose-binding as the two main substitutions (Y307Q and G127M) possibly responsible for the much lower affinity of CebE^reti^ for cellotriose are not conserved in CebE proteins of strains belonging to species *S. turgidiscabies* and *S. caniscabies*. Importantly, it has to be noted that our work also provides the structural basis for CebE-mediated uptake of cellobiose and cellotriose by saprophytic non-pathogenic *Streptomyces* species that actively participate in the mineralization of the plant decaying matter.

## Acknowledgments

The work of S.J. and B.D was supported by ‘Aspirant’ grants 1.A250.13 and 1.A618.18, from the ‘Fonds de la Recherche Scientifique’ (FNRS), respectively, a FRIA grant from the FNRS for N.S. (FRIA 1.E.116.21), and a FNRS grant “Crédit de recherche” (grant CDR/OL J.0158.21) to S.R. The work of F.K. is supported by a FNRS grant “Projet de recherche” (grant PDR/T.0121.22) and an “Action de Recherche Concertée” from Fédération Wallonie-Bruxelles (grant ARC 21/25-08). F.K. and S.R. are research and senior-research associates of the FRS-FNRS (Brussels, Belgium), respectively. We are very grateful for the assistance and support of the team of beamline PROXIMA 2A at the Soleil synchrotron.

---

**Figure S1.**
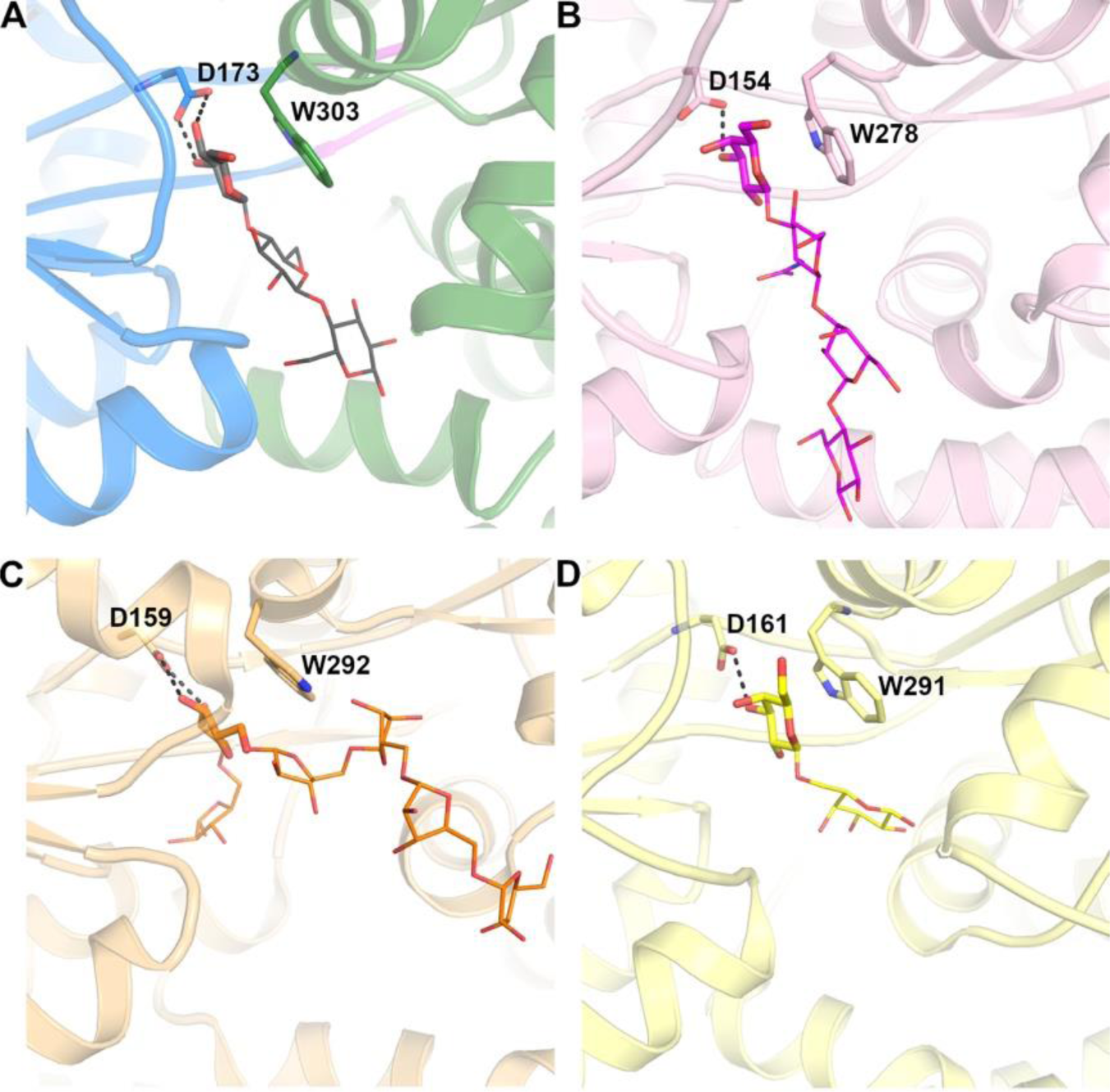
Comparison of saccharide binding mode in four SBP proteins. **A**. Cartoon representation of the CebE:cellotriose complex with coloring scheme as Figure 2. The aspartate and tryptophane conserved in SBP proteins are represented as sticks. **B**, **C** and **D**, similar representation for the GL-BP:lacto-N-tetraose, AbnE:arabinohexaose, and Bal6GBP:β-1,6-galactobiose complexes respectively.

## References

1. Liu H, Carvalhais LC, Crawford M, Singh E, Dennis PG, Pieterse CMJ, Schenk PM. 2017. Inner Plant Values: Diversity, Colonization and Benefits from Endophytic Bacteria. Front Microbiol 8.

2. Berlemont R, Martiny AC. 2013. Phylogenetic Distribution of Potential Cellulases in Bacteria. Appl Environ Microbiol 79:1545–1554.

3. Book AJ, Lewin GR, McDonald BR, Takasuka TE, Wendt-Pienkowski E, Doering DT, Suh S, Raffa KF, Fox BG, Currie CR. 2016. Evolution of High Cellulolytic Activity in Symbiotic Streptomyces through Selection of Expanded Gene Content and Coordinated Gene Expression. PLOS Biol 14:e1002475.

4. Book AJ, Lewin GR, McDonald BR, Takasuka TE, Doering DT, Adams AS, Blodgett JAV, Clardy J, Raffa KF, Fox BG, Currie CR. 2014. Cellulolytic Streptomyces Strains Associated with Herbivorous Insects Share a Phylogenetically Linked Capacity To Degrade Lignocellulose. Appl Environ Microbiol 80:4692–4701.

5. Saini A, Aggarwal NK, Sharma A, Yadav A. 2015. Actinomycetes: A Source of Lignocellulolytic Enzymes. Enzyme Res 2015:279381.

6. Lakhundi S, Siddiqui R, Khan NA. 2015. Cellulose degradation: a therapeutic strategy in the improved treatment of Acanthamoeba infections. Parasit Vectors 8:23.

7. Lim J-H, Lee C-R, Dhakshnamoorthy V, Park JS, Hong S-K. 2016. Molecular characterization of Streptomyces coelicolor A(3) SCO6548 as a cellulose 1,4-β-cellobiosidase. FEMS Microbiol Lett 363:fnv245.

8. Takasuka TE, Book AJ, Lewin GR, Currie CR, Fox BG. 2013. Aerobic deconstruction of cellulosic biomass by an insect-associated Streptomyces. Sci Rep 3:1030.

9. Jourdan S, Francis IM, Kim MJ, Salazar JJC, Planckaert S, Frère J-M, Matagne A, Kerff F, Devreese B, Loria R, Rigali S. 2016. The CebE/MsiK Transporter is a Doorway to the Cello-oligosaccharide-mediated Induction of Streptomyces scabies Pathogenicity. Sci Rep 6:27144.

10. Schlösser A, Jantos J, Hackmann K, Schrempf H. 1999. Characterization of the binding protein-dependent cellobiose and cellotriose transport system of the cellulose degrader Streptomyces reticuli. Appl Environ Microbiol 65:2636–2643.

11. Schlösser A, Kampers T, Schrempf H. 1997. The Streptomyces ATP-binding component MsiK assists in cellobiose and maltose transport. J Bacteriol 179:2092–2095.

12. Hurtubise Y, Shareck F, Kluepfel D, Morosoli R. 1995. A cellulase/xylanase-negative mutant of Streptomyces lividans 1326 defective in cellobiose and xylobiose uptake is mutated in a gene encoding a protein homologous to ATP-binding proteins. Mol Microbiol 17:367–377.

13. Deflandre B, Thiébaut N, Planckaert S, Jourdan S, Anderssen S, Hanikenne M, Devreese B, Francis I, Rigali S. 2020. Deletion of bglC triggers a genetic compensation response by awakening the expression of alternative beta-glucosidase. Biochim Biophys Acta BBA - Gene Regul Mech 1863:194615.

14. Jourdan S, Francis IM, Deflandre B, Tenconi E, Riley J, Planckaert S, Tocquin P, Martinet L, Devreese B, Loria R, Rigali S. 2018. Contribution of the β-glucosidase BglC to the onset of the pathogenic lifestyle of Streptomyces scabies. Mol Plant Pathol 19:1480–1490.

15. Deflandre B, Stulanovic N, Planckaert S, Anderssen S, Bonometti B, Karim L, Coppieters W, Devreese B, Rigali S 2022. The virulome of Streptomyces scabiei in response to cello-oligosaccharide elicitors. Microb Genomics 8:000760.

16. Marushima K, Ohnishi Y, Horinouchi S. 2009. CebR as a master regulator for cellulose/cellooligosaccharide catabolism affects morphological development in Streptomyces griseus. J Bacteriol 191:5930–5940.

17. Francis IM, Bergin D, Deflandre B, Gupta S, Salazar JJC, Villagrana R, Stulanovic N, Ribeiro Monteiro S, Kerff F, Loria R, Rigali S. 2023. Role of Alternative Elicitor Transporters in the Onset of Plant Host Colonization by Streptomyces scabiei 87–22. 2. Biology 12:234.

18. Johnson EG, Joshi MV, Gibson DM, Loria R. 2007. Cello-oligosaccharides released from host plants induce pathogenicity in scab-causing Streptomyces species. Physiol Mol Plant Pathol 71:18–25.

19. Joshi MV, Bignell DRD, Johnson EG, Sparks JP, Gibson DM, Loria R. 2007. The AraC/XylS regulator TxtR modulates thaxtomin biosynthesis and virulence in Streptomyces scabies. Mol Microbiol 66:633–642.

20. Planckaert S, Jourdan S, Francis IM, Deflandre B, Rigali S, Devreese B. 2018. Proteomic Response to Thaxtomin Phytotoxin Elicitor Cellobiose and to Deletion of Cellulose Utilization Regulator CebR in Streptomyces scabies. J Proteome Res 17:3837– 3852.

21. Liu J, Nothias L-F, Dorrestein PC, Tahlan K, Bignell DRD. 2021. Genomic and Metabolomic Analysis of the Potato Common Scab Pathogen Streptomyces scabiei. ACS Omega 6:11474–11487.

22. Haq IU, Mukhtar Z, Anwar-ul-Haq M, Liaqat S. 2023. Deciphering host–pathogen interaction during Streptomyces spp. infestation of potato. Arch Microbiol 205:222.

23. Schlösser A, Schrempf H. 1996. A Lipid-Anchored Binding Protein is a Component of an ATP-Dependent Cellobiose/Cellotriose-Transport System from the Cellulose Degrader Streptomyces reticuli. Eur J Biochem 242:332–338.

24. Jourdan S, Francis IM, Deflandre B, Loria R, Rigali S. 2017. Tracking the Subtle Mutations Driving Host Sensing by the Plant Pathogen Streptomyces scabies. mSphere 2:e00367–16.

25. Kabsch W. 2010. XDS. Acta Crystallogr D Biol Crystallogr 66:125–132.

26. Jumper J, Evans R, Pritzel A, Green T, Figurnov M, Ronneberger O, Tunyasuvunakool K, Bates R, Žídek A, Potapenko A, Bridgland A, Meyer C, Kohl SAA, Ballard AJ, Cowie A, Romera-Paredes B, Nikolov S, Jain R, Adler J, Back T, Petersen S, Reiman D, Clancy E, Zielinski M, Steinegger M, Pacholska M, Berghammer T, Bodenstein S, Silver D, Vinyals O, Senior AW, Kavukcuoglu K, Kohli P, Hassabis D. 2021. Highly accurate protein structure prediction with AlphaFold. 7873. Nature 596:583–589.

27. McCoy AJ, Grosse-Kunstleve RW, Adams PD, Winn MD, Storoni LC, Read RJ. 2007. Phaser crystallographic software. 4. J Appl Crystallogr 40:658–674.

28. Emsley P, Lohkamp B, Scott WG, Cowtan K. 2010. Features and development of Coot. 4. Acta Crystallogr D Biol Crystallogr 66:486–501.

29. Vagin AA, Steiner RA, Lebedev AA, Potterton L, McNicholas S, Long F, Murshudov GN. 2004. REFMAC5 dictionary: organization of prior chemical knowledge and guidelines for its use. Acta Crystallogr D Biol Crystallogr 60:2184–2195.

30. Hiard S, Marée R, Colson S, Hoskisson PA, Titgemeyer F, van Wezel GP, Joris B, Wehenkel L, Rigali S. 2007. PREDetector: a new tool to identify regulatory elements in bacterial genomes. Biochem Biophys Res Commun 357:861–864.

31. Rigali S, Nivelle R, Tocquin P. 2015. On the necessity and biological significance of threshold-free regulon prediction outputs. Mol Biosyst 11:333–337.

32. Berntsson RP-A, Smits SHJ, Schmitt L, Slotboom D-J, Poolman B. 2010. A structural classification of substrate-binding proteins. FEBS Lett 584:2606–2617.

33. Chandravanshi M, Tripathi SK, Kanaujia SP. 2021. An updated classification and mechanistic insights into ligand binding of the substrate-binding proteins. FEBS Lett 595:2395–2409.

34. Theilmann MC, Fredslund F, Svensson B, Leggio LL, Hachem MA. 2019. Substrate preference of an ABC importer corresponds to selective growth on β-(1,6)-galactosides in Bifidobacterium animalis subsp. lactis. J Biol Chem 294:11701–11711.

35. Lansky S, Salama R, Shulami S, Lavid N, Sen S, Schapiro I, Shoham Y, Shoham G. 2020. Carbohydrate-Binding Capability and Functional Conformational Changes of AbnE, an Arabino-oligosaccharide Binding Protein. J Mol Biol 432:2099–2120.

36. Suzuki R, Wada J, Katayama T, Fushinobu S, Wakagi T, Shoun H, Sugimoto H, Tanaka A, Kumagai H, Ashida H, Kitaoka M, Yamamoto K. 2008. Structural and Thermodynamic Analyses of Solute-binding Protein from Bifidobacterium longum Specific for Core 1 Disaccharide and Lacto-N-biose I *. J Biol Chem 283:13165–13173.

37. Chandravanshi M, Samanta R, Kanaujia SP. 2020. Conformational Trapping of a β-Glucosides-Binding Protein Unveils the Selective Two-Step Ligand-Binding Mechanism of ABC Importers. J Mol Biol 432:5711–5734.

38. Song Y, DiMaio F, Wang RY-R, Kim D, Miles C, Brunette TJ, Thompson J, Baker D. 2013. High-Resolution Comparative Modeling with RosettaCM. Structure 21:1735– 1742.

39. Guan D, Grau BL, Clark CA, Taylor CM, Loria R, Pettis GS. 2012. Evidence that thaxtomin C is a pathogenicity determinant of Streptomyces ipomoeae, the causative agent of Streptomyces soil rot disease of sweet potato. Mol Plant-Microbe Interact MPMI 25:393–401.

40. King RR, Lawrence CH, Clark MC, Calhoun LA. 1989. Isolation and characterization of phytotoxins associated with Streptomyces scabies. J Chem Soc Chem Commun 849– 850.

41. Doeven MK, van den Bogaart G, Krasnikov V, Poolman B. 2008. Probing Receptor-Translocator Interactions in the Oligopeptide ABC Transporter by Fluorescence Correlation Spectroscopy. Biophys J 94:3956–3965.

